# Sen1 is required for RNA Polymerase III transcription termination in an R-loop independent manner

**DOI:** 10.1101/571737

**Authors:** Julieta Rivosecchi, Marc Larochelle, Camille Teste, Frédéric Grenier, Amélie Malapert, Emiliano P. Ricci, Pascal Bernard, François Bachand, Vincent Vanoosthuyse

**Affiliations:** Laboratoire de Biologie et Modélisation de la Cellule, Université de Lyon, CNRS UMR 5239, Ecole Normale Supérieure de Lyon, Université Claude Bernard Lyon 1, 46 allée d’Italie F-69364 Lyon, France.; Département de Biochimie, Université de Sherbrooke, Sherbrooke, QC J1E4K8, Canada; Centre de Recherche du CHUS, Université de Sherbrooke, Sherbrooke, QC J1H5N4, Canada

**Keywords:** R-loops, RNA Polymerase III, Senataxin, Transcription termination

## Abstract

R-loop disassembly by the human helicase Senataxin contributes to genome stability and to proper transcription termination at a subset of RNA polymerase II genes. Whether Senataxin-mediated R-loop disassembly also contributes to transcription termination at other classes of genes has remained unclear. Here we show in fission yeast that Senataxin^Sen1^ promotes efficient termination of RNA Polymerase III (RNAP3) transcription *in vivo*. In the absence of Senataxin^Sen1^, RNAP3 accumulates downstream of the primary terminator at RNAP3-transcribed genes and produces long exosome-sensitive 3’-extended transcripts. Importantly, neither of these defects was affected by the removal of R-loops. The finding that Senataxin^Sen1^ acts as an ancillary factor for RNAP3 transcription termination *in vivo* challenges the pre-existing view that RNAP3 terminates transcription autonomously. We propose that Senataxin is a cofactor for transcription termination that has been co-opted by different RNA polymerases in the course of evolution.

## INTRODUCTION

Senataxin is a conserved DNA/RNA helicase whose deficiency has been implicated in the neurological disorders amyotrophic lateral sclerosis type 4 (ALS4) and ataxia-ocular apraxia type 2 (AOA2). How different Senataxin mutations contribute to the development of diseases with distinct pathologies remains unclear (Groh et al., 2017). Concordant observations in human cells and budding yeast have established that Senataxin is important for transcription termination of at least a subset of RNAP2-transcribed genes (Porrua and Libri, 2013; Skourti-Stathaki et al., 2011; Steinmetz et al., 2006), although the mechanisms involved probably differ in both species as budding yeast Senataxin^Sen1^ contributes to RNAP2 transcription termination as part of the Nrd1-Nab3-Sen1 (NNS) complex, which is not conserved in human cells. In addition, Senataxin has been implicated in the repair of DNA damage (Andrews et al., 2018; Cohen et al., 2018; Li et al., 2016) and in the resolution of transcription-replication conflicts (Alzu et al., 2012; Richard et al., 2013; Yüce and West, 2013). Both budding and fission yeast homologues of Senataxin can translocate in a 5’ to 3’ direction on either single-stranded DNA or RNA *in vitro* (Han et al., 2017; Kim et al., 1999; Martin-Tumasz and Brow, 2015) and it is believed that long, co-transcriptional DNA:RNA hybrids (also known as R-loops) represent a critical substrate of Senataxin *in vivo* (reviewed in Groh et al., 2017). Current models propose that the stabilization of R-loops upon Senataxin inactivation underlies the associated transcription termination and DNA repair defects. This proposal however has been somewhat challenged by the observation that budding yeast Senataxin^Sen1^ could directly dissociate pre-assembled RNAP2 transcription elongation complexes *in vitro* by translocating on the nascent RNA, even in the absence of DNA:RNA hybrids (Han et al., 2017; Porrua and Libri, 2013), suggesting that Senataxin has potentially R-loop-independent functions in the control of transcription.

The fission yeast *Schizosaccharomyces pombe* expresses two non-essential homologues of Senataxin, Senataxin^Sen1^ and Senataxin^Dbl8^. Surprisingly, transcription termination at RNAP2-transcribed genes is largely unaffected by lack of either or both homologues (Larochelle et al., 2018; Lemay et al., 2016) and to date, the roles of the fission yeast Senataxin enzymes have remained largely unknown. We reported previously that fission yeast Senataxin^Sen1^ associates with RNAP3 and is recruited to specific tRNA genes (Legros et al., 2014) but its function at RNAP3-transcribed genes has remained unclear. RNAP3 is predominantly implicated in the transcription of the short and abundant tRNA and 5S rRNA species. Internal promoter sequences called A- and B-box recruit the TFIIIC complex, which helps to position TFIIIB upstream of the Transcription Start Site (TSS). TFIIIB in turn recruits the 17 subunits RNAP3 complex at the TSS to initiate transcription (reviewed in (Schramm and Hernandez, 2002)). In fission yeast, an upstream TATA box assists TFIIIC in recruiting TFIIIB and is essential for the proper recruitment of RNAP3 (Hamada et al., 2001). *In vitro* transcription assays have indicated that once loaded, RNAP3 can terminate transcription autonomously upon reaching a transcription termination signal, which is constituted by a simple stretch of five thymine residues on the non-template strand (Mishra and Maraia, 2019), although this number may vary for different genes and organisms (reviewed in (Arimbasseri et al., 2013)). The C37/53/11 sub-complex of RNAP3 is particularly important for this intrinsic transcription termination mechanism (Arimbasseri et al., 2013). Interestingly, however, low levels of read-through transcription are observed at many tRNA genes *in vivo* in budding yeast, especially at tRNA genes with weaker terminator sequences (Turowski et al., 2016). The resulting 3’-extended transcripts are degraded by specific mechanisms involving the RNA exosome and the poly(A)-binding protein Nab2 (Turowski et al., 2016). In addition, human RNAP3 was also found in the 3’ regions of many tRNA genes (Orioli et al., 2011), suggesting that in distantly related systems, RNAP3 frequently overrides primary termination signals. These observations suggest that efficient RNAP3 transcription termination might be more challenging *in vivo* than suggested by *in vitro* studies. However, to what extent robust RNAP3 transcription termination requires the support of ancillary factors *in vivo* remains elusive.

We previously showed that unstable R-loops form at tRNA genes in fission yeast (Legros et al., 2014). Similar observations have been made in budding yeast (El Hage et al., 2014), humans (Chen et al., 2017) and plants (Xu et al., 2017), suggesting that R-loop formation is a conserved feature of RNAP3 transcription. The genome-wide stabilization of R-loops by the deletion of RNase H had a mild impact on pre-tRNA processing in budding yeast (El Hage et al., 2014). However, whether these mild perturbations were a direct consequence of R-loop stabilization in *cis* at tRNA genes was not addressed. Thus, the contribution of R-loops to RNAP3 transcription still remains largely unknown.

Our previous work revealed that fission yeast tRNA genes form R-loops and recruit Senataxin^Sen1^. As both R-loops and Senataxin were previously proposed to facilitate transcription termination of RNAP2 in humans (Skourti-Stathaki et al., 2011), we investigated the possibility that Senataxin^Sen1^ together with R-loops might participate in transcription termination of fission yeast RNAP3. We find that Senataxin^Sen1^ associates primarily with RNAP3 transcription units at the genome-wide level and does indeed promote proper RNAP3 transcription termination, albeit in a manner that is insensitive to the presence of R-loops. Thus, in contrast to conclusions drawn from *in vitro* studies, our work reveals the need for a protein cofactor to ensure robust RNAP3 transcription termination *in vivo*.

## RESULTS

### Sen1 associates primarily with RNAP3-transcribed genes

We reported previously that fission yeast Senataxin^Sen1^ (thereafter referred to as Sen1) associates with specific tRNA genes in an R-loop independent manner (Legros et al., 2014). However, the extent of Sen1 recruitment to all RNAP3-transcribed genes (tRNA, 5S rRNA, *srp7* and U6 snRNA) had remained unclear. We therefore used chromatin immunoprecipitation assays coupled to high-throughput sequencing (ChIP-seq) to analyse the genome-wide distribution of Sen1 relative to RNAP2 and RNAP3 occupancy. Analysis of ChIP-seq data confirmed that Sen1 is primarily enriched at tRNA and 5S rRNA genes, which also showed specific and robust RNAP3 binding (Fig 1A). In contrast, RNAP2-transcribed genes, as exemplified by the strongly expressed *tef3* gene in Figure 1A, showed only background Sen1 signal. Sen1 was also enriched at the RNAP3-transcribed *snu6* (Fig 1B) and *srp7* (Fig 1C) loci. A breakdown of all aligned ChIP-seq reads from two independent experiments revealed that more than 85% of Sen1-associated regions correspond to RNAP3-transcribed genes (Fig 1D, tRNA and 5S rRNA). This contrasts to Seb1 (Lemay et al., 2016), the fission yeast homologue of the NNS component Nrd1, that associates primarily with RNAP2-transcribed genes (Fig 1D). This is consistent with previous observations that Sen1 and Seb1 do not form a functional complex in fission yeast (Larochelle et al., 2018; Legros et al., 2014; Lemay et al., 2016). Overall, we observed a positive genome-wide correlation between ChIP-seq signals of Sen1 and two independent subunits of RNAP3 but not with RNAP2 (Fig 1E). We confirmed the enrichment of Sen1 at RNAP3-transcribed sites using ChIP-qPCR (Fig 1F). Interestingly, chromosome-organizing clamps (COC) sites, which recruit TFIIIC but not RNAP3 (Noma et al., 2006), were not enriched for Sen1 (Fig 1F). This is consistent with our observation that Sen1 physically associates with RNAP3 but not with TFIIIC (Legros et al., 2014) and suggests that the RNAP3 transcription complex might recruit Sen1 to target genes. To test this possibility, we mutated the upstream TATA box of a model tRNA gene, *SPCTRNAARG.10* (tRNA^ARG^_UCG_), in order to interfere with the recruitment of RNAP3 specifically at this locus. The mutated TATA-less *SPCTRNAARG.10* locus showed reduced levels of both RNAP3 and Sen1 (Fig 1G), indicating that the TATA box-dependent recruitment of RNAP3 is important for Sen1 binding. Collectively, these data indicate that the main targets of fission yeast Sen1 correspond to RNAP3-transcribed genes.

**Figure 1.**
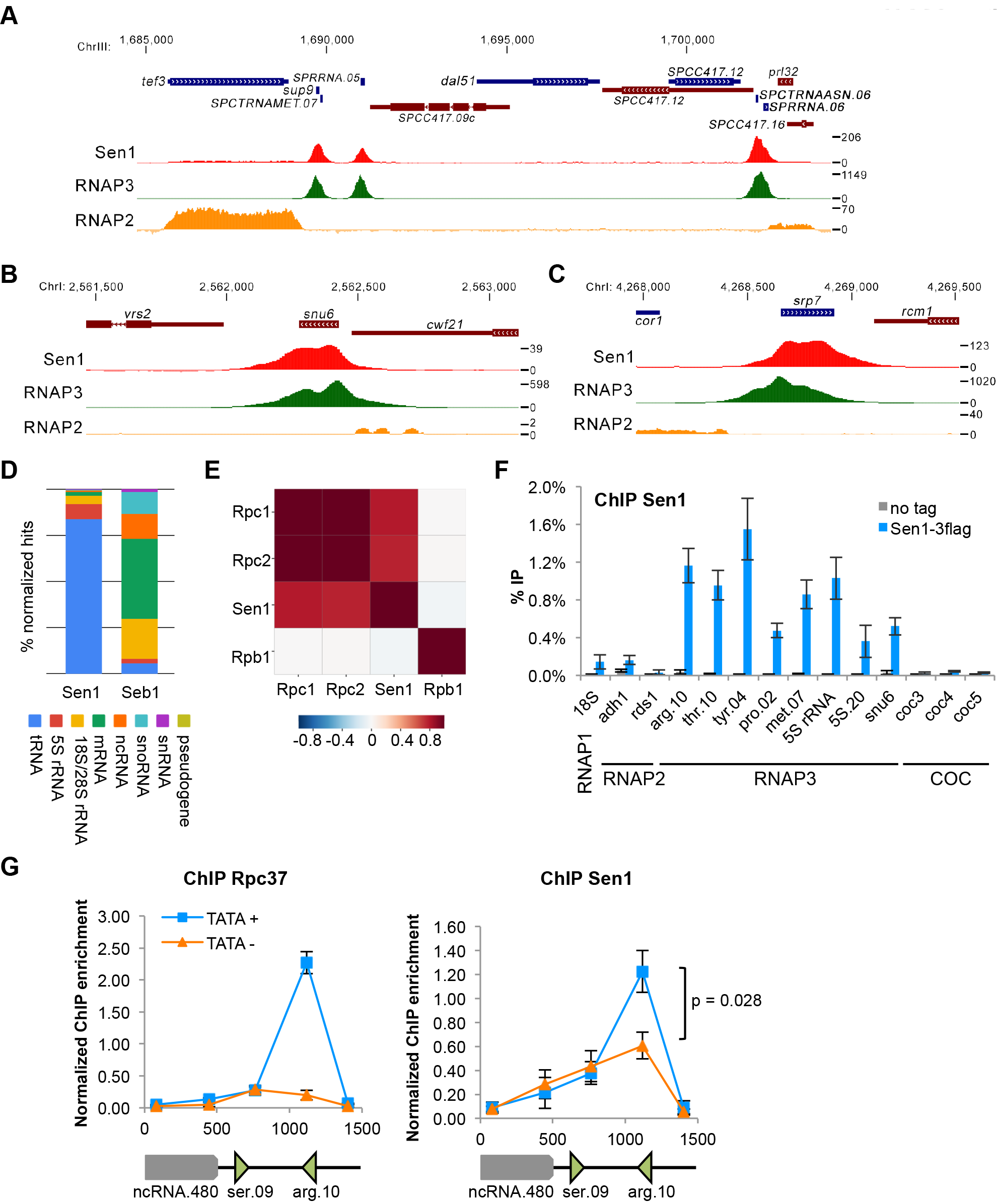
Sen1 associates predominantly with RNAP3-transcribed genes. **(ABC)** Snapshots of ChIP-seq signals of Sen1 (this study), RNAP3 (this study) and RNAP2 (data from (Larochelle et al., 2018)) across **(A)** a 25 kb region of chromosome 3 containing several tRNA and 5S rRNA genes, **(B)** the *snu6* and **(C)** the *srp7* loci. **(D)** Comparison of the distribution of ChIP-seq reads across the indicated categories of genes for Sen1 (this study) and Seb1 (data from (Lemay et al., 2016)). **(E)** Genome-wide pairwise Pearson correlation coefficient matrix at a resolution of 10 bp. **(F)** ChIP-qPCR analysis of Flag-tagged Sen1 at the indicated loci in a population of cycling cells (mean ± standard deviation from 4 biological replicates). **(G)** ChIP-qPCR analysis of the Flag-tagged RNAP3 subunit Rpc37 and Flag-tagged Sen1 across the *SPCTRNAARG.10* tRNA locus, whose TATA box was either mutated (TATA−) or not (TATA+). The enrichment values were normalized to *SPCTRNATHR.10* (mean ± standard deviation from 4 biological replicates; p-value obtained using the Wilcoxon-Mann Whitney statistical test).

### Sen1 is required for normal RNAP3 transcription

Next we investigated whether lack of Sen1 affects RNAP3 transcription. Using ChIP-qPCR, we detected increased amount of RNAP3 at all sites tested in the absence of Sen1 (Fig 2A). Strikingly, this accumulation of RNAP3 over its target genes was not associated with an increased amount of RNAP3 transcripts in the cell (Fig 2B). The steady state level of 5S RNAs remained unchanged and the overall levels of tRNA detected in the absence of Sen1 were even slightly reduced (Fig 2B). This reduction was more apparent when Dis3-mediated tRNA degradation (Gudipati et al., 2012; Schneider et al., 2012) was impaired (Fig 2B, compare lanes 3-4). Since the steady state levels of tRNAs is controlled by the equilibrium between synthesis and Dis3-mediated degradation (Gudipati et al., 2012; Schneider et al., 2012), to deplete Dis3 to reduce activity of the nuclear exosome is expected to uncover the impact of Sen1 on RNAP3 transcription. The small reduction in tRNA levels in the single *sen1*Δ mutant was confirmed using gene-specific RT-qPCR (Fig 2C). Taken together, these results indicate that lack of Sen1 interferes with RNAP3 transcription.

**Figure 2:**
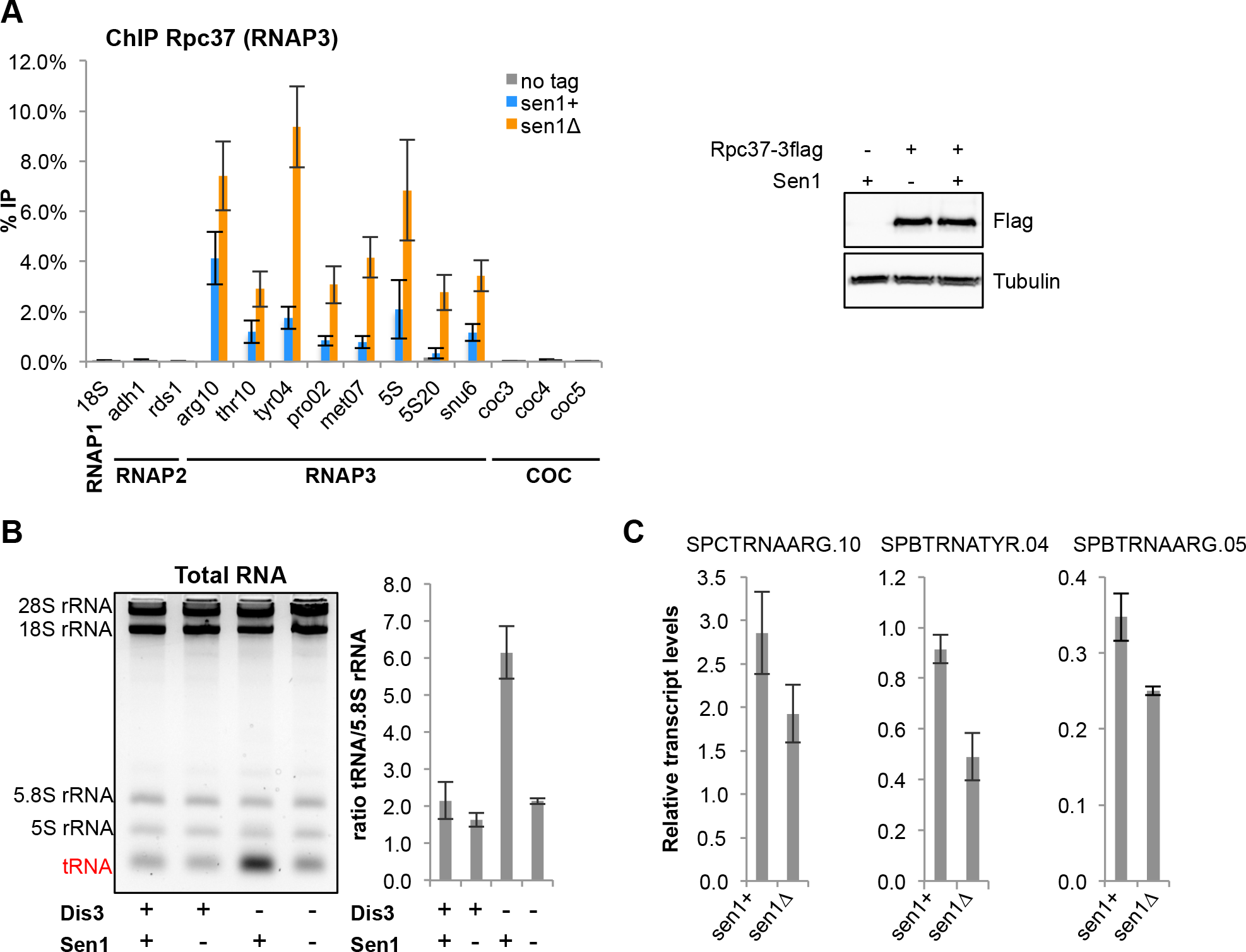
Sen1 is required for normal RNAP3 transcription. **(A)** (left) ChIP-qPCR analysis of Rpc37 in the presence or absence of Sen1 at the indicated loci in a population of cycling cells (mean ± standard deviation from 4 biological replicates). (right) Western blot analysis of Rpc37 protein levels in the presence or absence of Sen1. Tubulin was used as a loading control. **(B)** (left) Total RNA from the indicated strains were separated on a 2,8% agarose gel. (right) Quantification of overall tRNA levels in 3 independent experiments. **(C)** Strand-specific RT-qPCR was used to quantify the indicated RNAP3 transcripts. Transcript levels were normalized to *act1* (mean ± standard deviation from 3 biological replicates).

### The accumulation of RNAP3 on chromatin in the absence of Sen1 is independent of R-loops

Sen1 is believed to antagonize R-loop formation and R-loops were shown to interfere with transcription elongation, at least when they form close to the TSS (Belotserkovskii et al., 2017). We have shown previously that RNase H-sensitive R-loops form at tRNA genes in fission yeast (Hartono et al., 2018; Legros et al., 2014). To test whether the stabilization of R-loops at tRNA genes could underlie the accumulation of RNAP3 in the absence of Sen1, we expressed RNase H1 from *Escherichia coli* (RnhA) under the control of the strong *nmt1* promoter in fission yeast cells. We showed previously that this strategy was sufficient to remove R-loops at tRNA genes (Hartono et al., 2018; Legros et al., 2014). Using both dot-blots with the S9.6 antibody (Boguslawski et al., 1986) to quantify overall DNA:RNA hybrid levels and R-ChIP to monitor R-loop formation at tRNA genes (Legros et al., 2014), we found that lack of Sen1 did not noticeably change the amount of R-loops (Fig 3AB). In addition, we confirmed that RnhA expression was sufficient to completely remove R-loops in the absence of Sen1 (Fig 3AB). However, this treatment did not alter the accumulation of RNAP3, indicating that R-loops do not contribute to the accumulation of RNAP3 in the *sen1*Δ mutant (Fig 3C). Conversely, stabilization of R-loops at tRNA genes by the deletion of both endogenous RNase H1 and RNase H2 (*rnh1*Δ*rnh201*Δ) (Legros et al., 2014) did not result in the accumulation of RNAP3 (Fig 3C). These results therefore establish that the stabilization of R-loops does not account for the accumulation of RNAP3 in Sen1-deficient cells.

**Figure 3:**
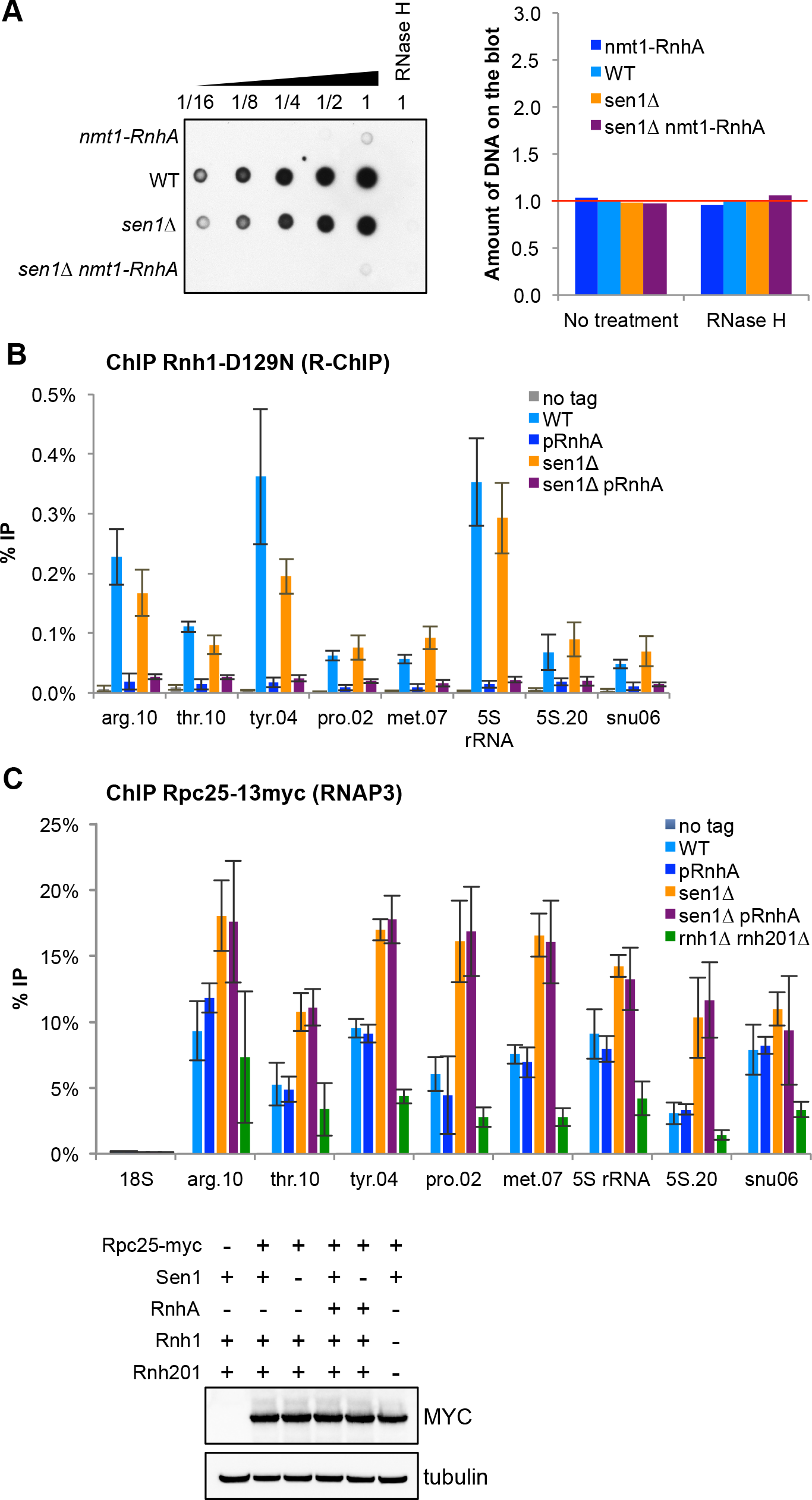
Sen1 regulates RNAP3 transcription in an R-loop independent manner. Cells were grown in minimal medium during 18 hours to induce the strong expression of RnhA. **(A)** (left) Genomic DNA from the indicated strains was spotted on a membrane and its levels of DNA:RNA hybrids were quantified using the S9.6 antibody. RNase H-treated genomic DNA was used as a specificity control. (right) The amount of genomic DNA spotted was quantified by qPCR. **(B)** R-ChIP using a catalytically-inactive RNase H1 (Rnh1-D129N) was used to quantify R-loop formation at RNAP3-transcribed genes (mean ± standard deviation from 4 biological replicates). **(C)** (top) ChIP-qPCR of the 13myc-tagged RNAP3 subunit Rpc25 in the indicated strains at the indicated loci (mean ± standard deviation of 4 biological replicates). (bottom) Western blot analysis of Rpc25 protein levels in the indicated strains. Tubulin was used as a loading control.

### Sen1 is required for RNAP3 transcription termination

We next analysed the effect of a Sen1 deficiency on the genome-wide distribution of RNAP3 by comparing ChIP-seq profiles of Rpc1 and Rpc2 in sen1+ and *sen1*Δ strains. Strikingly, in the absence of Sen1, the distribution of both Rpc1 and Rpc2 displayed increased density downstream of most tRNA and 5S rRNA genes (as exemplified on Fig 4A&B), as well as at *srp7* (Fig 4C), consistent with read-through transcription by RNAP3. Evidence of delayed transcription termination in the *sen1*Δ mutant was also noted at the *snu6* gene, albeit at more modest levels (Fig EV1). Importantly, averaging Rpc1 and Rpc2 ChIP-seq signals over all isolated tRNA and 5S rRNA genes confirmed that the distribution pattern of RNAP3 is globally extended at the 3’ end in the absence of Sen1 (Fig 4D). ChIP followed by qPCR analysis at several candidate loci confirmed that the domain occupied by RNAP3 was wider in the absence of Sen1 and that RNAP3 accumulated downstream of its natural transcription termination sites (Fig 4E). Notably, this accumulation downstream of the transcription termination site was not altered upon RnhA expression, confirming that it did not result from the stabilization of R-loops (Fig EV2). Together, these results show that Sen1 is required for RNAP3 termination at the genome-wide level in an R-loop independent manner.

**Figure 4:**
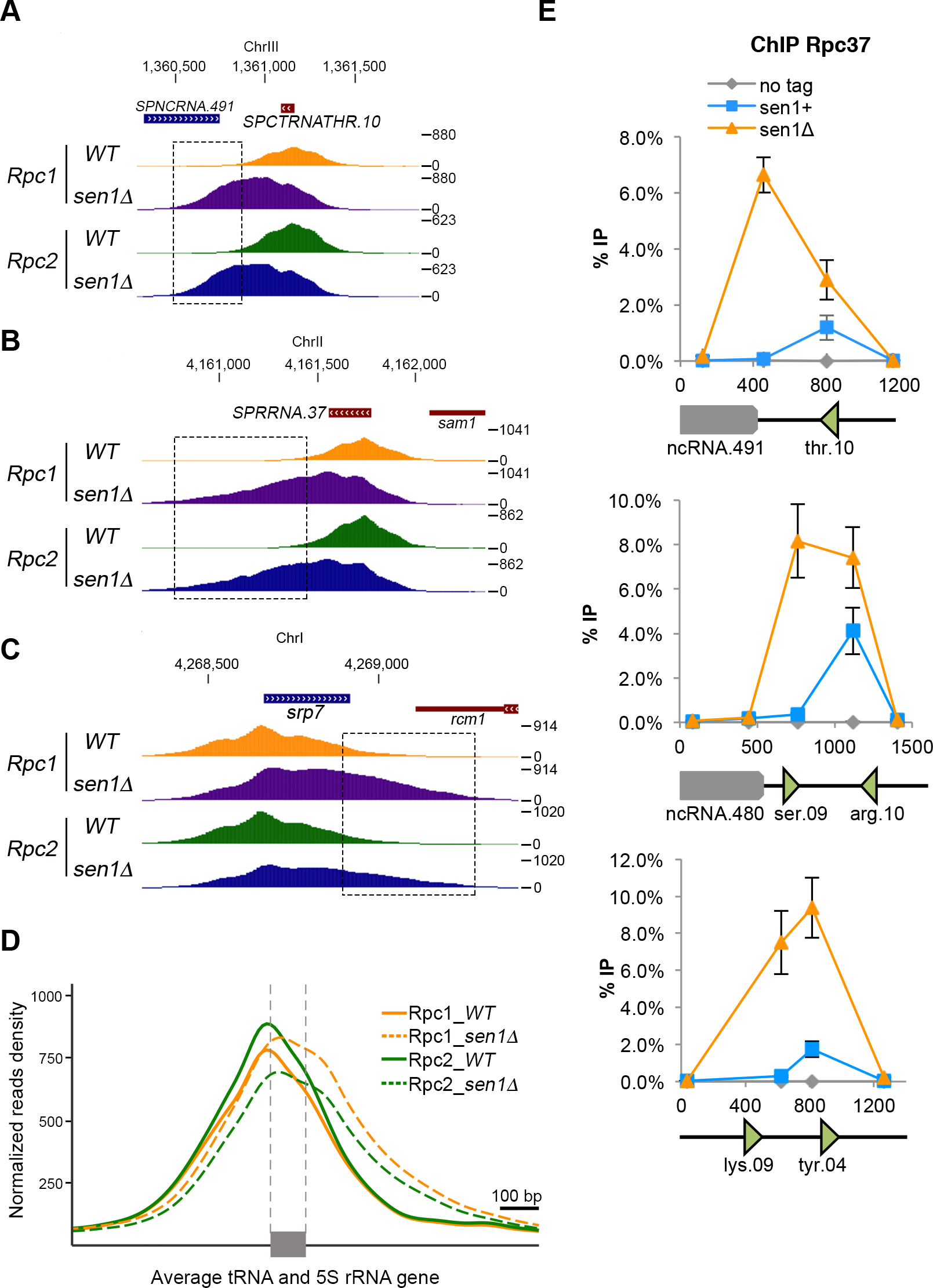
Sen1 is required for RNAP3 transcription termination. **(ABC)** Snapshots of ChIP-seq signals of the RNAP3 subunits Rpc1 and Rpc2 in the presence or absence of Sen1 across a representative **(A)** tRNA gene, **(B)** 5S rRNA gene and **(C)** *srp7*. Boxed regions highlight the increased density of reads in the downstream region of genes in the absence of Sen1. **(D)** Average ChIP-seq profile of Rpc1 and Rpc2 across all isolated tRNA and 5S rRNA genes in the presence and absence of Sen1. **(E)** Scanning of Rpc37-3flag occupancy at three different tDNA loci in the absence of Sen1 by ChIP-qPCR (mean ± standard deviation from 6 biological replicates).

### Read-through tRNA transcripts accumulate in the absence of Sen1

We used several independent assays to demonstrate that the transcription termination defects associated with lack of Sen1 resulted in the production of 3’-extended transcripts. First, we used a genetic assay that translates a transcription termination defect at the synthetic tRNA DRT5T construct into a change of colour of the yeast colonies from red to white (Iben et al., 2011) (Fig 5A, see scheme of the construct at the top). Briefly, a transcription termination defect allows the synthesis of a suppressor tRNA that suppresses the accumulation of a red pigment caused by the *ade6-704* mutation, resulting in white colonies in limiting adenine conditions. As a positive control in this assay, we mutated the valine residue at position 189 in the Rpc37 subunit of RNAP3 into an aspartate residue (*rpc37-V189D*), as overexpression of this mutant was shown to interfere with transcription termination in a dominant-negative manner (Rijal and Maraia, 2013). Here we mutated instead the endogenous *rpc37* gene and established that the *rpc37-V189D* mutant is viable but displays transcription termination defects (Fig 5A). Similarly, in the absence of Sen1 but not in the absence of its close homologue Dbl8, colonies turned white in the presence of the DRT5T construct, indicating that Sen1 but not Dbl8 is required for robust transcription termination at DRT5T (Fig 5A). Consistent with the idea that Sen1 contributes to transcription termination of RNAP3-transcribed genes, we found that Sen1 becomes essential for cell viability when the termination is impaired by the *rpc37-V189D* mutant (Fig EV3).

**Figure 5:**
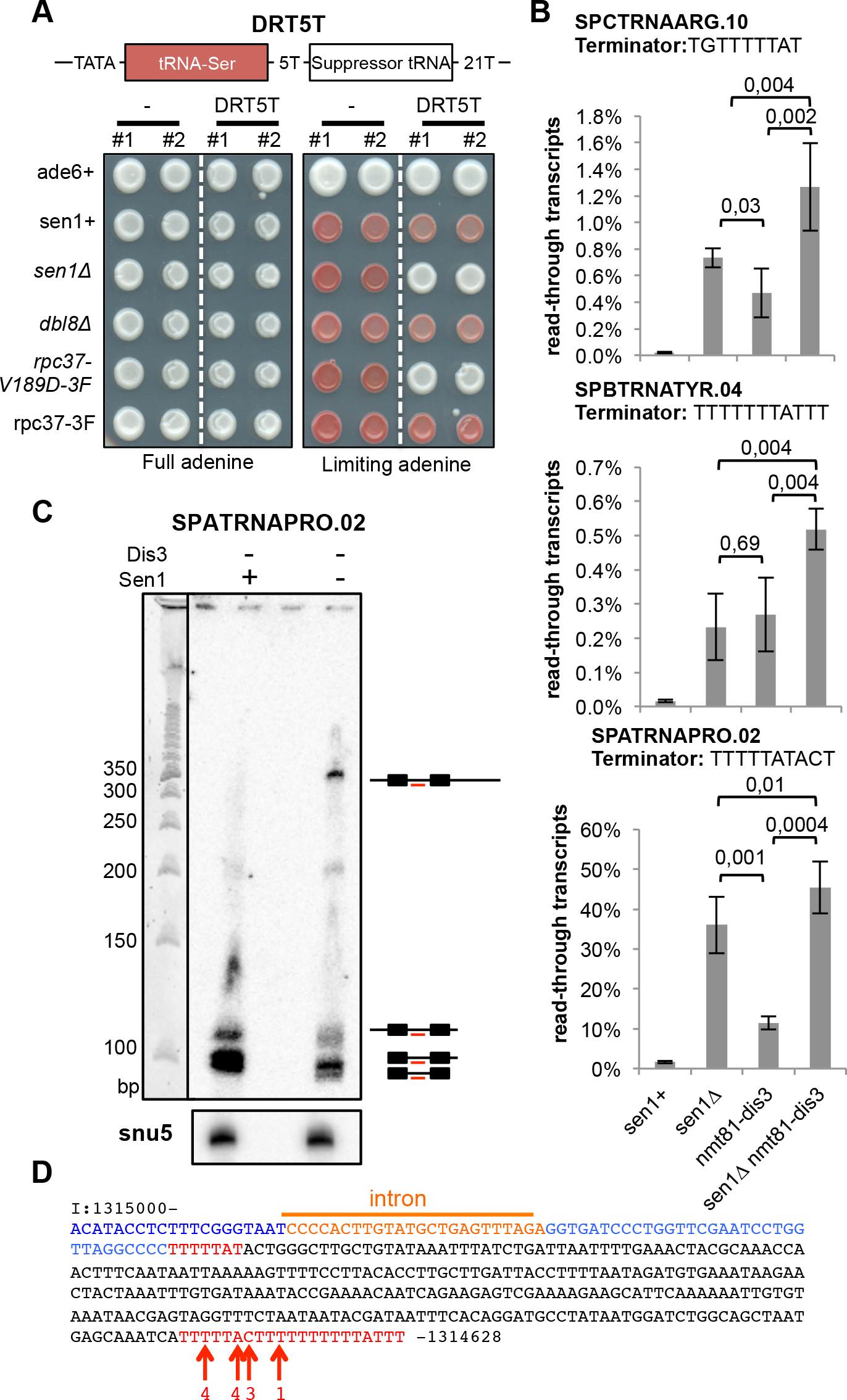
Lack of Sen1 produces extended read-through tRNA transcripts. **(A)** Cells of the indicated genotypes that carried or not the DRT5T dimeric tRNA construct (schematized on top) were grown either in the presence of the optimum concentration of adenine (left) or in the presence of a limiting concentration of adenine (right). Two independent clones of the same genotype (#1 and #2) were used. See text for details. 3F refers to the 3 Flag epitope tag at the C-terminus of Rpc37. **(B)** Strand-specific RT-qPCR was used to quantify the levels of read-through transcripts (see Methods). The mean ± standard deviation from 4 biological replicates is represented here. P-values were obtained using the Wilcoxon-Mann Whitney statistical test. **(C)** Northern blot analysis of the tRNA *SPATRNAPRO.02* using an intron-specific probe (TCTAAACTCAGCATACAAGTGGGG). U5 snRNA was used as a loading control. **(D)** Sequence of the ~350 nt-long read-through transcript at *SPATRNAPRO.02*. Residues in blue represent the sequence of the mature tRNA. Residues in red represent potential terminator sequences. Red arrows show the 3’ end nucleotide of the read-through transcripts. The numbers indicate the number of times the sequenced transcripts terminated at the indicated position.

To confirm the production of read-through transcripts at endogenous tRNA genes in the absence of Sen1, we used strand-specific RT-qPCR. Using this approach, we detected read-through transcripts in the absence of Sen1 but not Dbl8 at several tRNA genes (Fig EV4). To rule out that those read-through transcripts only resulted from the defective degradation of naturally occurring longer transcripts (Turowski et al., 2016), we quantified these read-through transcripts in the absence of the RNA exosome subunit Dis3. If Sen1 was only involved in the RNA exosome-dependent degradation of naturally-occurring read-through transcripts, the amount of read-through transcripts found in RNA exosome mutants should not change upon deletion of Sen1. As shown in Fig 5B, we found that the amount of read-through transcripts detected in RNA exosome mutants increased significantly in the absence of Sen1, strongly suggesting that those longer transcripts did not result from an impaired RNA exosome activity. Using Northern blots, we detected a predominant ~350 nt-long extended transcript at the intron-containing *SPATRNAPRO.02* (tRNA^PRO^_CGG_) in the absence of Sen1 (Fig 5C). Sequencing of the 3’ end of this transcript confirmed that it included a 3’ extension and showed that it terminated at a strong distal terminator sequence (TTTTTACTTTTTTTTTTATTT) located 274 bp downstream of the primary terminator (Fig 5D), showing that consistent with our ChIP-seq data RNAP3 continues transcribing downstream of the primary terminator sequence in the absence of Sen1. Finally, we showed that the accumulation of read-through transcripts in the absence of Sen1 was unchanged after RnhA expression, reinforcing the idea that read-through transcription in the absence of Sen1 was not a consequence of R-loop stabilization (Fig EV5). Taken together, our data indicate that Sen1 prevents the synthesis of long, aberrant read-through tRNAs by promoting efficient termination of RNAP3.

### A strong terminator sequence compensates for lack of Sen1

It was shown previously that a stretch of 21 thymine residues is sufficient to provide robust transcription termination even in the presence of a termination-defective RNAP3 (Iben et al., 2011; Rijal and Maraia, 2013) and our analysis of read-through transcripts at *SPATRNAPRO.02* (Fig 5C&D) was consistent with the idea that a strong terminator sequence can overcome the requirement for Sen1 for RNAP3 transcription termination. We therefore tested whether strengthening the primary terminator sequence could rescue the defects observed in the absence of Sen1. To do this, we replaced the endogenous terminator sequence of *SPCTRNAARG.10* by a stretch of 23 thymine residues (Fig 6A). Strikingly, the presence of such a strong terminator sequence was sufficient to suppress both the accumulation of RNAP3 downstream of the terminator (Fig 6B) and the accumulation of read-through transcripts in the absence of Sen1 (Fig 6C). Note however that the presence of this super-terminator at *SPCTRNAARG.10* did not affect the production of read-through transcripts at the neighbouring *SPCTRNASER.09* (Fig 6C). Similarly, the introduction of a terminator composed of 20 consecutive thymine residues at *SPCTRNATHR.10* was sufficient to suppress the accumulation of RNAP3 downstream of the terminator (Fig EV6). Altogether, these observations demonstrate that the RNAP3 molecules that accumulate downstream of the terminator sequence in the absence of Sen1 correspond to RNAP3 molecules that overrode the terminator sequence and conclusively establish that Sen1 is required for robust transcription termination of RNAP3-transcribed genes.

**Figure 6.**
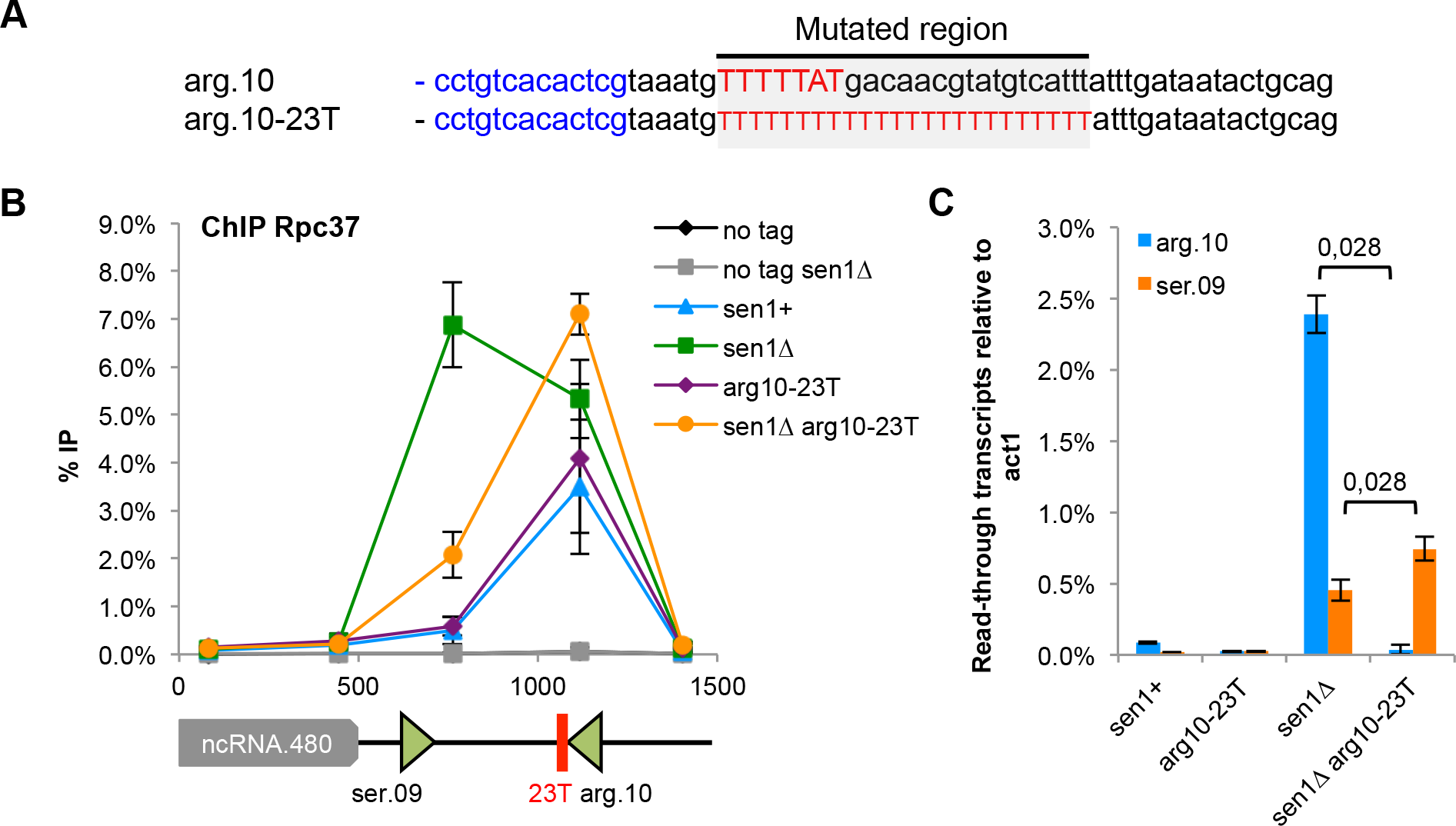
A strong terminator sequence compensates for lack of Sen1. **(A)** Sequence of the engineered strong terminator (*arg10-23T*) at the *SPCTRNAARG.10* gene. **(B)** ChIP-qPCR analysis of Rpc37 around *SPCTRNAARG.10* gene in the strong terminator mutant (mean ± standard deviation from 4 biological replicates). **(C)** Strand specific RT-qPCR was used to quantify read-through transcripts at *SPCTRNAARG.10* (arg10) and *SPCTRNASER.09* (ser09) in the strong terminator mutant (mean ± standard deviation from 4 biological replicates; p-value obtained using the Wilcoxon-Mann Whitney statistical test).

## DISCUSSION

RNAP3 transcribes the abundant structural tRNA and 5S rRNA transcripts. In contrast to RNAP2, whose transcription termination relies on the coordinated assembly of dedicated protein complexes at the 3’ of genes, termination of RNAP3 is generally thought to rely on the autonomous destabilization of the elongation complex upon reaching dedicated terminator DNA sequences. Here we challenge this notion by showing that the highly conserved DNA/RNA helicase Sen1 is a cofactor of RNAP3 required for global termination of RNAP3 transcription in fission yeast.

### Senataxin homologues are conserved regulators of transcription termination

The function of Senataxin homologues in the termination of transcription at specific RNAP2-transcribed genes is well established in budding yeast and in mammals (Porrua and Libri, 2013; Skourti-Stathaki et al., 2011) but was previously ruled out for the fission yeast homologues Senataxin^Sen1^ and Senataxin^Dbl8^ (Larochelle et al., 2018; Lemay et al., 2016). Here we show that the function of Senataxin^Sen1^ in transcription termination is conserved in fission yeast, albeit for another RNA polymerase. This suggests that a contribution to transcription termination is an ancestral function of Senataxin homologues and it is tempting to speculate that their mode of action in promoting transcription termination is conserved for RNAP3.

In budding yeast, Senataxin^Sen1^ was shown to interact with all three RNA polymerases (Yüce and West, 2013), opening the possibility that it might contribute to transcription termination of all three RNA polymerases. Consistent with this, it was proposed previously that Senataxin^Sen1^ contributes also to RNAP1 transcription termination in budding yeast (Kawauchi et al., 2008). Interestingly, our unpublished observations indicate that the other Senataxin homologue in fission yeast, Senataxin^Dbl8^, interacts with RNAP1, opening the possibility that it might assist RNAP1 transcription termination. Whether or not another DNA&RNA helicase assists RNAP2 transcription termination in fission yeast in a Senataxin-like manner remains to be established. Similarly, the interaction between mammalian Senataxin and RNAP3 has not been reported and it is unclear at this stage whether Senataxin or another DNA&RNA helicase contributes to RNAP3 transcription termination in mammals.

### Fission yeast Sen1 promotes robust RNAP3 elongation and termination *in vivo*

*In vitro* studies have led to the conclusion that RNAP3 could terminate transcription autonomously, without the need for accessory factors (reviewed in (Arimbasseri et al., 2013)). In contrast to this view, our data clearly show that Senataxin^Sen1^ is required for efficient termination of RNAP3 transcription *in vivo*. This suggests that *in vitro* transcription assays, although extremely informative, do not fully recapitulate RNAP3 transcription. *In vitro* transcription assays do not yet recapitulate the chromatin environment and often rely on the artificial assembly of an elongating RNAP3 without the need for TFIIIB or TFIIIC. It is possible that the role of Sen1 in transcription termination becomes dispensable in these conditions. This could indicate that the way RNAP3 is loaded onto the DNA template has an impact on the way transcription terminates, as suggested previously (Wang and Roeder, 1998). Alternatively, it is possible that transcription elongation rates are higher *in vivo* than *in vitro* and demand a more efficient Senataxin^Sen1^-dependent transcription termination mechanism, in line with what was proposed for Senataxin^Sen1^-mediated RNAP2 termination in budding yeast (Hazelbaker et al., 2013).

Our set of ChIP experiments has shown that RNAP3 levels increased at RNAP3-transcribed genes in the absence of Senataxin^Sen1^. Interestingly, this accumulation was not accompanied by an increase in the amount of mature tRNA. On the contrary, we measured reduced amount of tRNA in the absence of Sen1 when Dis3-dependent tRNA degradation was impaired (Fig 2B), suggesting that nascent tRNA production is reduced. We therefore speculate that the accumulation of RNAP3 that was detected in the absence of Senataxin^Sen1^ reflects transcription elongation defects rather than enhanced transcription rates. Importantly, as the introduction of a super-terminator sequence at *SPACTRNAARG.10* suppressed the transcription termination defects in *sen1Δ* cells, but did not reduce RNAP3 back to normal levels on this tRNA gene (Fig 6B), this suggests that the putative transcription elongation defects are independent from the transcription termination defects observed in Sen1-deficient cells.

### Could a Mfd-like release and catch-up mechanism explain the different roles of Sen1?

We still ignore the precise mode of action of fission yeast Senataxin^Sen1^ in RNAP3 transcription termination. Recently, it was proposed that the Mfd helicase in *Escherichia coli* could both nudge forward weakly paused RNAP and induce the dissociation of stalled RNAP using a release and catch-up mechanism (Le et al., 2018). We speculate that such a release and catch-up mechanism could underlie the roles of fission yeast Senataxin^Sen1^ in RNAP3 transcription: through translocation, Senataxin^Sen1^ could nudge forward RNAP3 molecules that are weakly paused at gene-internal pause sites such as TFIIIC-binding sites (Turowski et al., 2016) and thereby facilitate transcription elongation, or release RNAP3 molecules that are stalled at canonical terminator sequences to promote transcription termination. Mfd was shown previously to translocate autonomously on DNA, whilst both budding yeast and fission yeast Senataxin^Sen1^ were shown to translocate on both DNA and RNA, albeit at greater rate on DNA (Han et al., 2017; Kim et al., 1999; Martin-Tumasz and Brow, 2015). However, translocation on the nascent RNA was sufficient to explain the ability of budding yeast Senataxin^Sen1^ to dissociate elongation complexes *in vitro* (Han et al., 2017).

Importantly, the release and catch-up mechanism was also proposed to underlie the role of Mfd in both transcription-coupled repair and transcription-replication conflict resolution (Le et al., 2018). Strikingly, budding yeast Senataxin^Sen1^ has also been implicated in both transcription-coupled repair (Li et al., 2016) and transcription-replication conflict resolution (Alzu et al., 2012; Brambati et al., 2018; Mischo et al., 2011), strengthening the analogy with Mfd. A unifying way of interpreting the different roles of Senataxin^Sen1^ could therefore be to propose that it is targeted to chromatin through its interaction with RNA polymerases and that it subsequently patrols chromatin locally to facilitate the resolution of R-loops and/or stalled elongation complexes through a release and catch-up mechanism. The identity of the RNA polymerase involved would change in different organisms or for different paralogues in the same organism. In this scenario, Senataxin^Sen1^ would not recognize R-loops directly. Consistent with this, Senataxin was not identified as an R-loop binding protein in two recent proteomic screens (Cristini et al., 2018; Wang et al., 2018). We propose instead that the ancestral function of Senataxin is to dissociate stalled transcription elongation complexes from chromatin.

## MATERIALS AND METHODS

### Fission yeast strains and culture

The list of all the strains used in this study is given in Supplemental_Table_S1. Standard genetic crosses were employed to construct all strains. *sen1-3flag*, and *rpc37-3flag* were generated using a standard PCR procedure. Cells were grown at 30°C in complete YES+adenine medium or in synthetic PMG medium as indicated. The expression of the *dis3* gene driven by the *nmt81* promoter was repressed by the addition of 60 μM of thiamine to the PMG medium. To induce the expression of *E. coli* RnhA from by the *nmt1* promoter, cells were grown in PMG minimal medium lacking thiamine for a minimum of 18h.

### Mutagenesis of Rpc37

To generate *rpc37-V189D-3Flag*, the 3’ end of *rpc37* was first amplified by PCR and cloned into pCR Blunt II-TOPO using the Zero Blunt II TOPO PCR Cloning Kit (Invitrogen Life Technologies). Site-directed PCR mutagenesis was then carried out to mutate the codon corresponding to valine 189 (GTC into GAC) using the QuikChange II Site-Directed Mutagenesis Kit (Agilent Technologies). Overlapping PCR was used to add a C-terminus 3Flag epitope and a marker conferring resistance to Nourseothricin and the resulting PCR fragment was transformed into fission yeast using routine protocols. Proper integration of the mutation at the endogenous locus was verified by PCR and sequencing.

### Mutagenesis of the primary terminator of *SPCTRNAARG.10*

The TATA-less and super-terminator mutants of *SPCTRNAARG.10* (*arg10 TATA-less* and *arg10-23T*) were first synthesized (GeneCust Europe). The mutagenized *SPCTRNAARG.10* gene was then transformed into fission yeast and its correct integration was selected by counter-selecting on FOA the loss of the *ura4* gene previously integrated at *SPCTRNAARG.10.* The correct integration of the mutation was confirmed by sequencing.

### Chromatin Immunoprecipitation (ChIP)

1,5.10^8^ cells were cross-linked with 1% formaldehyde (Sigma) at 18°C for 30 minutes. After 3 washes with cold PBS, the cells were frozen in liquid nitrogen. Frozen cells were then lysed in cold lysis buffer (Hepes-KOH 50mM [pH 7.5], NaCl 140mM, EDTA 1mM, Triton 1%, Na-deoxycholate 0.1%, PMSF Phenylmethanesulfonyl fluoride 1 mM) with glass beads using a Precellys 24 mill (Bertin Technology). To fragment the chromatin, the lysates were sonicated at 4°C using a Covaris S220 or Diagenode Bioruptor sonicator. Immuno-precipitation was done overnight at 4°C using Protein A-coupled or Protein G-coupled Dynabeads previously incubated with anti-GFP A11122 antibody (Invitrogen), anti-Flag antibody (M2 Sigma) and anti-Myc 9E10 (Sigma). The immunoprecipitated complexes were washed for 5’ successively with: Wash I buffer (20mM Tris pH 8, 150 mM NaCl, 2mM EDTA, 1% Triton-X100, 0.1% SDS), Wash II buffer (20mM Tris pH 8, 500mM NaCl, 2 mM EDTA, 1% Triton-X100, 0.1% SDS) and Wash III buffer (20mM Tris pH 8, 1mM EDTA, 0.5% Na-deoxycholate, 1% Igepal, 250mM LiCl). After two additional washes in Tris EDTA pH 8, the beads were resuspended in 10% Chelex resin (Biorad) and incubated at 98°C for 10’. After addition of 2 μL of 10 mg/mL proteinase K, the mixture was incubated at 43°C for 1 hour, then for another 10’ at 98°C. After centrifugation, the supernatant was collected and analyzed by qPCR in a thermocycler Rotor Gene (Qiagen) using the primers listed in Supplemental_Table_S2.

### Western Blotting

Protein extraction was performed using the TCA (trichloroacetic acid)-glass beads method. 10^8^ cells were centrifuged 3’ at 3400 rpm, resuspended in 20% TCA and then lysed using glass beads in a Precellys 24 mill (Bertin Technology). After centrifugation (4’ at 13 krpm), the resulting pellets were resuspended in sample buffer (0,1M Tris-HCl pH 9.5, 20% Glycerol, 4% SDS (sodium dodecyl sulphate), 0.2% bromophenol blue, 715mM β-mercaptoethanol), incubated 5’ at 100°C and centrifugated again at 13 krpm for 4’. The resulting supernatants were separated using SDS-PAGE on 7,5 % polyacrylamide gels and transferred onto nitrocellulose using a semi-dry transfer system. Anti-Myc (A-14 Santa Cruz Biotechnology) and anti-Flag (M2 Sigma) antibodies were used for immunodetection of proteins and revealed using ECL-based reagents. An anti-tubulin antibody (TAT1), courtesy of Prof. Keith Gull (Oxford) was used as loading control.

### RNA techniques

Total RNA was extracted from logarithmical growing cells (2.10^8^) by the standard hot-phenol method. The remaining traces of genomic DNA digest were digested with DNAse I (Ambion) and the integrity of RNAs was verified by electrophoresis on 0,8% agarose gels. Total RNA was reverse-transcribed using SuperScript III (Invitrogen) according to the manufacturer’s instructions using the strand-specific primers listed in Supplemental_Table_S2. To quantify read-through transcripts at specific tRNA genes, two independent RT reactions were carried out in parallel: RT1 used a priming oligonucleotide placed downstream of the primary terminator and RT2 used a priming oligonucleotide placed in the gene body upstream of the primary terminator. For both RT1 and RT2, an *act1*-specific priming oligonucleotide was also used as internal control. The resulting cDNAs were quantified by quantitative PCR (qPCR) using a Rotor Gene machine (Qiagen) and primers specific for *act1* and the gene body of the tRNA of interest. The proportion of read-through transcripts was expressed using the ratio RT1/RT2 (read-through tRNA transcripts/total tRNA transcripts). For Northern blots, 10 μg of total RNAs were separated on 10% polyacrylamide–8M urea gels and transferred onto a nylon membrane (GE Healthcare Amersham Hybond −N+). The membrane was then UV cross-linked and dried at 80°C for 30 minutes. After incubation with Church Buffer for 30 minutes at 37°C, the membrane was hybridized overnight at 37°C with a ^32^P-labeled DNA oligo antisense to the intron of *SPATRNAPRO.02*. The blot was then washed 4 times with 1X SSC + 0.1% SDS and scanned using a Phosphorimager Typhoon FLA 9500 - GE Healthcare.

### Mapping of the 3’ end of read-through transcripts at *SPATRNAPRO.02*

20 μg of total RNA were separated on a 10% polyacrylamide–8M urea gel. The 200 to 600 bp-long RNAs were extracted from the gel and purified, before a preadenylated RNA adaptor was ligated in 3’ as described previously (Heyer et al., 2015). This 3’ adaptor was used for retro-transcription as described (Heyer et al., 2015) and *SPATRNAPRO.02*-derived cDNAs were amplified by PCR using the primer Pro.02 qL1 (5’-ACATACCTCTTTCGGGTAATCC-3’). The PCR fragment obtained was cloned into pCR Blunt II TOPO using the Zero Blunt II TOPO PCR Cloning Kit (Invitrogen Life Technologies) and sequenced.

### Transcription termination assay

Strains carrying the *ade6-704* mutation and the DRT5T dimeric construct were obtained from the Maraia laboratory. Standard genetic crosses were employed to introduce these reporter constructs in the strains of interest. At least two independent strains for each genotype were then plated on YES medium depleted or not of adenine for 3 days at 30°C.

#### Library preparation and Illumina sequencing

DNA librairies for ChIP-seq experiments were prepared as described previously (Lemay et al., 2016) using the SPARK DNA Sample Prep Kit Illumina Platform (Enzymatics) according to the manufacturer’s instructions.

#### ChIP-seq processing

Briefly, the raw reads were trimmed using Trimmomatic version 0.32 (Bolger et al., 2014) with parameters ILLUMINACLIP:2:30:15 LEADING:30 TRAILING:30 MINLEN:23, and quality inspection was conducted using FastQC version 0.11.4 (https://www.bioinformatics.babraham.ac.uk/projects/fastqc/). The trimmed reads from all data sets were aligned using BWA version 0.7.12-r1039 (Li and Durbin, 2010) with the algorithm mem and the parameter -split onto the sequence of the *S. pombe* ASM294v2. Note that no filtering on mapQ was performed in order to avoid discarding the signal at regions of the genome that are duplicated (BWA is randomly assigning the reads), but only primary alignments were kept and we generated mappability tracks for various read length to help the interpretation of particular regions. Signal density files in BedGraph format were then generated using BEDTools genomecov version 2.17 (Quinlan and Hall, 2010) with default parameters, then converted in uniform 10 nt bins WIG files for further normalization steps (inspired by the script bedgraph_to_wig.py [https://gist.github.com/svigneau/8846527]).

#### ChIP-seq data analysis

Each signal density file was scaled based on sample’s sequencing depth, then the signal of the input data set was subtracted from its corresponding IP data set. The normalized WIG files were then encoded in bigWig format using the Kent utilities (Rhead et al., 2010). Visual inspection of the data was performed using an AssemblyHub on the UCSC Genome Browser (Casper et al., 2018).

The Versatile Aggregate Profiler (VAP) tool (Brunelle et al., 2015; Coulombe et al., 2014) version 1.1.0 was used to generate the average profile using the following parameters: annotation mode, absolute analysis method, 10 bp windows size, mean aggregate value, smoothing of 6, and missing data were considered as “0”. 131 isolated tRNA and 35 isolated 5S rRNA were used for the VAP analyses. We used the 3’ one reference point mode to generate the graph values and identified the average 5’ start using the 5’ one reference point mode (both graphs overlapped perfectly for all curves). Genome-wide Pearson correlation coefficients were calculated using the epiGeEC tool version 1.0 (Laperle et al., 2018).

To generate the % normalized hits distribution (Fig 1D), we used summarizeOverlaps from the R package GenomicAlignments (Lawrence et al., 2013) with the Union mode to count the aligned reads overlapping with each gene, both with the input and IP data. The counts were then normalized based on sample’s sequencing depth. Then the normalized input count was subtracted from the normalized IP count. Finally, the resulting counts were further normalized based on gene length. The % normalized hits for every type of gene were computed as the sum of the positive counts for a gene type divided by the sum of all positive count.

#### Code availability

All scripts used for data processing and statistical analysis were written in Python, Perl, or R, and are available upon request.

## ACKNOWLEDGMENTS

We are very grateful to Richard Maraia for sending reagents. We thank Thomas Diot for helping with the genome annotations of tRNA and their terminators in fission yeast. We thank the members of the LBMC biocomputing center for their advice, in particular Laurent Modolo and Hélène Polvèche. We are grateful to Domenico Libri and Odil Porrua for helpful discussions. This work was supported by a “Chaire d’Excellence” awarded to VV by the Agence Nationale pour la Recherche (ANR, Project TRACC, CHX11), by a “Projet Pluri-Equipe” (PPE 2016–2018) awarded by la Ligue contre le Cancer, comité du Rhône to VV and by the PRCE project “GeneSilencingByCondensin” (ANR-15-CE12-0002-01) and a “Projet Fondation ARC” (PJA 20151203343) awarded to PB by the Agence Nationale pour la Recherche (ANR) and the Fondation ARC pour la recherche sur le cancer, respectively. Work in the Bachand laboratory was supported by funding from the Natural Sciences and Engineering Research Council of Canada (NSERC) to F.B. (RGPIN-2017-05482). F.B. holds a Canada Research Chair in Quality Control of Gene Expression.

## AUTHOR CONTRIBUTIONS

VV conceived the study with input from PB, JR and FB. JR, CT, ER, AM and VV carried out all experiments except for the ChIP-seq experiments. ML, FG and FB carried out and analysed the ChIP-seq experiments. VV, JR, FB and PB discussed and interpreted the results. VV wrote the manuscript with input from FB and PB and all authors revised it.

## CONFLICT OF INTEREST

The authors declare that they have no conflict of interest

## EXPANDED VIEW FIGURE LEGENDS

**Figure EV1. Lack of Sen1 results in modest transcription termination defects at *snu6***. Snapshots of ChIP-seq signals of the RNAP3 subunits Rpc1 and Rpc2 in the presence or absence of Sen1 across the *snu6* locus. Boxed region highlights the increased density of reads downstream of *snu6* in the absence of Sen1.

**Figure EV2. Loss of R-loops upon expression of RnhA does not impact the distribution of RNAP3 in the absence of Sen1.** ChIP-qPCR analysis of the RNAP3 sub-unit Rpc25 in the indicated genotypes and at the indicated loci in a population of cycling cells (mean ± standard deviation from 2 biological replicates).

**Figure EV3. Sen1 function is essential in the termination-defective RNAP3 mutant *rpc37-V189D***. Tetrad dissection was used to show that the double mutant *sen1*Δ *rpc37-V189D* is dead.

**Figure EV4. Lack of Sen1 but not lack of its close homologue Dbl8 results in read-through transcripts at tRNA genes.** Strand-specific RT-qPCR was used to quantify the levels of read-through transcripts (see Methods). The mean ± standard deviation from 2 biological replicates is represented here.

**Figure EV5. RnhA expression does not change the proportion of read-through transcripts in the absence of Sen1.** Strand-specific RT-qPCR was used to quantify the levels of read-through transcripts (see Methods). The mean ± standard deviation from 3 biological replicates is represented here.

**Figure EV6. A super-terminator at *SPCTRNATHR.10* suppresses the accumulation of RNAP3 in 3’ when Sen1 is missing.** (A) Sequences of the engineered strong (thr10-20T) at the SPCTRNATHR.10 gene. (B) ChIP-qPCR analysis of Rpc37 around SPCTRNATHR.10 gene in the strong terminator mutant. (mean ± standard deviation from 4 biological replicates). The RNAP3-transcribed *SPBTRNAARG.04* (arg.04) and the RNAP1-transcribed rDNA (18S) were used as specificity controls.

## REFERENCES

Alzu, A., Bermejo, R., Begnis, M., Lucca, C., Piccini, D., Carotenuto, W., Saponaro, M., Brambati, A., Cocito, A., Foiani, M., et al. (2012). Senataxin associates with replication forks to protect fork integrity across RNA-polymerase-II-transcribed genes. Cell 151, 835–846.

Andrews, A.M., McCartney, H.J., Errington, T.M., D’Andrea, A.D., and Macara, I.G. (2018). A senataxin-associated exonuclease SAN1 is required for resistance to DNA interstrand cross-links. Nat. Commun. 9, 2592.

Arimbasseri, A.G., Rijal, K., and Maraia, R.J. (2013). Transcription termination by the eukaryotic RNA polymerase III. Biochim. Biophys. Acta 1829, 318–330.

Belotserkovskii, B.P., Soo Shin, J.H., and Hanawalt, P.C. (2017). Strong transcription blockage mediated by R-loop formation within a G-rich homopurine-homopyrimidine sequence localized in the vicinity of the promoter. Nucleic Acids Res. 45, 6589–6599.

Boguslawski, S.J., Smith, D.E., Michalak, M.A., Mickelson, K.E., Yehle, C.O., Patterson, W.L., and Carrico, R.J. (1986). Characterization of monoclonal antibody to DNA.RNA and its application to immunodetection of hybrids. J. Immunol. Methods 89, 123–130.

Bolger, A.M., Lohse, M., and Usadel, B. (2014). Trimmomatic: a flexible trimmer for Illumina sequence data. Bioinforma. Oxf. Engl. 30, 2114–2120.

Brambati, A., Zardoni, L., Achar, Y.J., Piccini, D., Galanti, L., Colosio, A., Foiani, M., and Liberi, G. (2018). Dormant origins and fork protection mechanisms rescue sister forks arrested by transcription. Nucleic Acids Res. 46, 1227–1239.

Brunelle, M., Coulombe, C., Poitras, C., Robert, M.-A., Markovits, A.N., Robert, F., and Jacques, P.-É. (2015). Aggregate and Heatmap Representations of Genome-Wide Localization Data Using VAP, a Versatile Aggregate Profiler. Methods Mol. Biol. Clifton NJ 1334, 273–298.

Casper, J., Zweig, A.S., Villarreal, C., Tyner, C., Speir, M.L., Rosenbloom, K.R., Raney, B.J., Lee, C.M., Lee, B.T., Karolchik, D., et al. (2018). The UCSC Genome Browser database: 2018 update. Nucleic Acids Res. 46, D762–D769.

Chen, L., Chen, J.-Y., Zhang, X., Gu, Y., Xiao, R., Shao, C., Tang, P., Qian, H., Luo, D., Li, H., et al. (2017). R-ChIP Using Inactive RNase H Reveals Dynamic Coupling of R-loops with Transcriptional Pausing at Gene Promoters. Mol. Cell 68, 745–757.e5.

Cohen, S., Puget, N., Lin, Y.-L., Clouaire, T., Aguirrebengoa, M., Rocher, V., Pasero, P., Canitrot, Y., and Legube, G. (2018). Senataxin resolves RNA:DNA hybrids forming at DNA double-strand breaks to prevent translocations. Nat. Commun. 9, 533.

Coulombe, C., Poitras, C., Nordell-Markovits, A., Brunelle, M., Lavoie, M.-A., Robert, F., and Jacques, P.-É. (2014). VAP: a versatile aggregate profiler for efficient genome-wide data representation and discovery. Nucleic Acids Res. 42, W485–493.

Cristini, A., Groh, M., Kristiansen, M.S., and Gromak, N. (2018). RNA/DNA Hybrid Interactome Identifies DXH9 as a Molecular Player in Transcriptional Termination and R-Loop-Associated DNA Damage. Cell Rep. 23, 1891–1905.

El Hage, A., Webb, S., Kerr, A., and Tollervey, D. (2014). Genome-wide distribution of RNA-DNA hybrids identifies RNase H targets in tRNA genes, retrotransposons and mitochondria. PLoS Genet. 10, e1004716.

Groh, M., Albulescu, L.O., Cristini, A., and Gromak, N. (2017). Senataxin: Genome Guardian at the Interface of Transcription and Neurodegeneration. J. Mol. Biol. 429, 3181–3195.

Gudipati, R.K., Xu, Z., Lebreton, A., Séraphin, B., Steinmetz, L.M., Jacquier, A., and Libri, D. (2012). Extensive degradation of RNA precursors by the exosome in wild-type cells. Mol. Cell 48, 409–421.

Hamada, M., Huang, Y., Lowe, T.M., and Maraia, R.J. (2001). Widespread use of TATA elements in the core promoters for RNA polymerases III, II, and I in fission yeast. Mol. Cell. Biol. 21, 6870–6881.

Han, Z., Libri, D., and Porrua, O. (2017). Biochemical characterization of the helicase Sen1 provides new insights into the mechanisms of non-coding transcription termination. Nucleic Acids Res. 45, 1355–1370.

Hartono, S.R., Malapert, A., Legros, P., Bernard, P., Chédin, F., and Vanoosthuyse, V. (2018). The Affinity of the S9.6 Antibody for Double-Stranded RNAs Impacts the Accurate Mapping of R-Loops in Fission Yeast. J. Mol. Biol. 430, 272–284.

Hazelbaker, D.Z., Marquardt, S., Wlotzka, W., and Buratowski, S. (2013). Kinetic competition between RNA Polymerase II and Sen1-dependent transcription termination. Mol. Cell 49, 55–66.

Heyer, E.E., Ozadam, H., Ricci, E.P., Cenik, C., and Moore, M.J. (2015). An optimized kit-free method for making strand-specific deep sequencing libraries from RNA fragments. Nucleic Acids Res. 43, e2.

Iben, J.R., Mazeika, J.K., Hasson, S., Rijal, K., Arimbasseri, A.G., Russo, A.N., and Maraia, R.J. (2011). Point mutations in the Rpb9-homologous domain of Rpc11 that impair transcription termination by RNA polymerase III. Nucleic Acids Res. 39, 6100–6113.

Kawauchi, J., Mischo, H., Braglia, P., Rondon, A., and Proudfoot, N.J. (2008). Budding yeast RNA polymerases I and II employ parallel mechanisms of transcriptional termination. Genes Dev. 22, 1082–1092.

Kim, H.D., Choe, J., and Seo, Y.S. (1999). The sen1(+) gene of Schizosaccharomyces pombe, a homologue of budding yeast SEN1, encodes an RNA and DNA helicase. Biochemistry (Mosc.) 38, 14697–14710.

Laperle, J., Hébert-Deschamps, S., Raby, J., Morais, D.A. de L., Barrette, M., Bujold, D., Bastin, C., Robert, M.-A., Nadeau, J.-F., Harel, M., et al. (2018). The epiGenomic Efficient Correlator (epiGeEC) tool allows fast comparison of user datasets with thousands of public epigenomic datasets. Bioinforma. Oxf. Engl.

Larochelle, M., Robert, M.-A., Hébert, J.-N., Liu, X., Matteau, D., Rodrigue, S., Tian, B., Jacques, P.-É., and Bachand, F. (2018). Common mechanism of transcription termination at coding and noncoding RNA genes in fission yeast. Nat. Commun. 9, 4364.

Lawrence, M., Huber, W., Pagès, H., Aboyoun, P., Carlson, M., Gentleman, R., Morgan, M.T., and Carey, V.J. (2013). Software for computing and annotating genomic ranges. PLoS Comput. Biol. 9, e1003118.

Le, T.T., Yang, Y., Tan, C., Suhanovsky, M.M., Fulbright, R.M., Inman, J.T., Li, M., Lee, J., Perelman, S., Roberts, J.W., et al. (2018). Mfd Dynamically Regulates Transcription via a Release and Catch-Up Mechanism. Cell 172, 344–357.e15.

Legros, P., Malapert, A., Niinuma, S., Bernard, P., and Vanoosthuyse, V. (2014). RNA processing factors Swd2.2 and Sen1 antagonize RNA Pol III-dependent transcription and the localization of condensin at Pol III genes. PLoS Genet. 10, e1004794.

Lemay, J.-F., Marguerat, S., Larochelle, M., Liu, X., van Nues, R., Hunyadkürti, J., Hoque, M., Tian, B., Granneman, S., Bähler, J., et al. (2016). The Nrd1-like protein Seb1 coordinates cotranscriptional 3’ end processing and polyadenylation site selection. Genes Dev. 30, 1558–1572.

Li, H., and Durbin, R. (2010). Fast and accurate long-read alignment with Burrows-Wheeler transform. Bioinforma. Oxf. Engl. 26, 589–595.

Li, W., Selvam, K., Rahman, S.A., and Li, S. (2016). Sen1, the yeast homolog of human senataxin, plays a more direct role than Rad26 in transcription coupled DNA repair. Nucleic Acids Res. 44, 6794–6802.

Martin-Tumasz, S., and Brow, D.A. (2015). Saccharomyces cerevisiae Sen1 Helicase Domain Exhibits 5’- to 3’-Helicase Activity with a Preference for Translocation on DNA Rather than RNA. J. Biol. Chem. 290, 22880–22889.

Mischo, H.E., Gómez-González, B., Grzechnik, P., Rondón, A.G., Wei, W., Steinmetz, L., Aguilera, A., and Proudfoot, N.J. (2011). Yeast Sen1 helicase protects the genome from transcription-associated instability. Mol. Cell 41, 21–32.

Mishra, S., and Maraia, R.J. (2019). RNA polymerase III subunits C37/53 modulate rU:dA hybrid 3’ end dynamics during transcription termination. Nucleic Acids Res. 47, 310–327.

Noma, K., Cam, H.P., Maraia, R.J., and Grewal, S.I.S. (2006). A role for TFIIIC transcription factor complex in genome organization. Cell 125, 859–872.

Orioli, A., Pascali, C., Quartararo, J., Diebel, K.W., Praz, V., Romascano, D., Percudani, R., van Dyk, L.F., Hernandez, N., Teichmann, M., et al. (2011). Widespread occurrence of non-canonical transcription termination by human RNA polymerase III. Nucleic Acids Res. 39, 5499–5512.

Porrua, O., and Libri, D. (2013). A bacterial-like mechanism for transcription termination by the Sen1p helicase in budding yeast. Nat. Struct. Mol. Biol. 20, 884–891.

Quinlan, A.R., and Hall, I.M. (2010). BEDTools: a flexible suite of utilities for comparing genomic features. Bioinforma. Oxf. Engl. 26, 841–842.

Rhead, B., Karolchik, D., Kuhn, R.M., Hinrichs, A.S., Zweig, A.S., Fujita, P.A., Diekhans, M., Smith, K.E., Rosenbloom, K.R., Raney, B.J., et al. (2010). The UCSC Genome Browser database: update 2010. Nucleic Acids Res. 38, D613–619.

Richard, P., Feng, S., and Manley, J.L. (2013). A SUMO-dependent interaction between Senataxin and the exosome, disrupted in the neurodegenerative disease AOA2, targets the exosome to sites of transcription-induced DNA damage. Genes Dev. 27, 2227–2232.

Rijal, K., and Maraia, R.J. (2013). RNA polymerase III mutants in TFIIFα-like C37 that cause terminator readthrough with no decrease in transcription output. Nucleic Acids Res. 41, 139–155.

Schneider, C., Kudla, G., Wlotzka, W., Tuck, A., and Tollervey, D. (2012). Transcriptome-wide analysis of exosome targets. Mol. Cell 48, 422–433.

Schramm, L., and Hernandez, N. (2002). Recruitment of RNA polymerase III to its target promoters. Genes Dev. 16, 2593–2620.

Skourti-Stathaki, K., Proudfoot, N.J., and Gromak, N. (2011). Human senataxin resolves RNA/DNA hybrids formed at transcriptional pause sites to promote Xrn2-dependent termination. Mol. Cell 42, 794–805.

Steinmetz, E.J., Warren, C.L., Kuehner, J.N., Panbehi, B., Ansari, A.Z., and Brow, D.A. (2006). Genome-wide distribution of yeast RNA polymerase II and its control by Sen1 helicase. Mol. Cell 24, 735–746.

Turowski, T.W., Leśniewska, E., Delan-Forino, C., Sayou, C., Boguta, M., and Tollervey, D. (2016). Global analysis of transcriptionally engaged yeast RNA polymerase III reveals extended tRNA transcripts. Genome Res. 26, 933–944.

Wang, Z., and Roeder, R.G. (1998). DNA topoisomerase I and PC4 can interact with human TFIIIC to promote both accurate termination and transcription reinitiation by RNA polymerase III. Mol. Cell 1, 749–757.

Wang, I.X., Grunseich, C., Fox, J., Burdick, J., Zhu, Z., Ravazian, N., Hafner, M., and Cheung, V.G. (2018). Human proteins that interact with RNA/DNA hybrids. Genome Res. 28, 1405–1414.

Xu, W., Xu, H., Li, K., Fan, Y., Liu, Y., Yang, X., and Sun, Q. (2017). The R-loop is a common chromatin feature of the Arabidopsis genome. Nat. Plants 3, 704–714.

Yüce, Ö., and West, S.C. (2013). Senataxin, defective in the neurodegenerative disorder ataxia with oculomotor apraxia 2, lies at the interface of transcription and the DNA damage response. Mol. Cell. Biol. 33, 406–417.

